# An insertion unique to SARS-CoV-2 exhibits superantigenic character strengthened by recent mutations

**DOI:** 10.1101/2020.05.21.109272

**Authors:** Mary Hongying Cheng, She Zhang, Rebecca A. Porritt, Moshe Arditi, Ivet Bahar

**Affiliations:** Department of Computational and Systems Biology, School of Medicine, University of Pittsburgh, Pittsburgh, PA 15261; Department of Pediatrics, Division of Pediatric Infectious Diseases and Immunology, Cedars-Sinai Medical Center, Los Angeles, CA 90048.; Department of Biomedical Sciences, Infectious and Immunologic Diseases Research Center, Cedars-Sinai Medical Center, Los Angeles, CA 90048.

**Keywords:** Covid-19, Superantigen, SARS-CoV-2 Spike, Toxic shock syndrome, Cytokine storm

## Abstract

Multisystem Inflammatory Syndrome in Children (MIS-C) associated with Coronavirus Disease 2019 (COVID-19) is a newly recognized condition in which children with recent SARS-CoV-2 infection present with a constellation of symptoms including hypotension, multiorgan involvement, and elevated inflammatory markers. These symptoms and the associated laboratory values strongly resemble toxic shock syndrome, an escalation of the cytotoxic adaptive immune response triggered upon the binding of pathogenic superantigens to MHCII molecules and T cell receptors (TCRs). Here, we used structure-based computational models to demonstrate that the SARS-CoV-2 spike (S) exhibits a high-affinity motif for binding TCR, interacting closely with both the α- and β-chains variable domains’ complementarity-determining regions. The binding epitope on S harbors a sequence motif unique to SARS-CoV-2 (not present in any other SARS coronavirus), which is highly similar in both sequence and structure to bacterial superantigens. Further examination revealed that this interaction between the virus and human T cells is strengthened in the context of a recently reported rare mutation (D839Y/N/E) from a European strain of SARS-CoV-2. Furthermore, the interfacial region includes selected residues from a motif shared between the SARS viruses from the 2003 and 2019 pandemics, which has intracellular adhesion molecule (ICAM)-like character. These data suggest that the SARS-CoV-2 S may act as a superantigen to drive the development of MIS-C as well as cytokine storm in adult COVID-19 patients, with important implications for the development of therapeutic approaches.

**Significance:** Although children have been largely spared from severe COVID-19 disease, a rare hyperinflammatory syndrome has been described in Europe and the East Coast of the United States, termed Multisystem Inflammatory Syndrome in Children (MISC). The symptoms and diagnostic lab values of MIS-C resemble those of toxic shock, typically caused by pathogenic superantigens stimulating excessive activation of the adaptive immune system. We show that SARS-CoV-2 spike has a sequence and structure motif highly similar to those of bacterial superantigens, and may directly bind to the T cell receptors. This sequence motif, not present in other coronaviruses, may explain the unique potential for SARS-CoV-2 to cause both MIS-C and the cytokine storm observed in adult COVID-19 patients.

---

**S**evere acute respiratory syndrome coronavirus 2 (SARS-CoV-2), which causes COVID-19, is a coronavirus closely related to SARS-CoV and Middle East Respiratory Syndrome (MERS) coronaviruses (1). COVID-19 can manifest in adults as a severe interstitial pneumonia with hyperinflammation while severe respiratory manifestations are rare in children (2–4). Recently, however, multisystem inflammatory system in children (MIS-C) has been recognized in patients that either tested positive for COVID-19 (by PCR or serology) or had epidemiological links to COVID-19 (5–7). After initial reports in UK (5), many cases have now been reported in Europe (6, 7), and New York (USA CDC). However, no such cases have been reported in China, Japan, or South Korea, which have also been severely impacted by the COVID-19 pandemic (ECDC).

MIS-C manifests as persistent fever and hyperinflammation with multi organ system involvement including cardiac, gastrointestinal, renal, hematologic, dermatologic and neurologic symptoms (5–7) which are highly reminiscent of toxic shock syndrome (TSS) (8, 9) (**Table 1**), rather than Kawasaki disease due to marked demographic, clinical, and laboratory differences (6). The similarities to TSS and the association of MIS-C with COVID-19 led us to hypothesize that SARS-CoV-2 may possess superantigenic fragments that induce an inflammatory cascade and perhaps also contribute to the hyperinflammation and cytokine storm features observed in severe adult COVID-19 cases (3, 4). The question we raised is: does SARS-CoV-2 S possess superantigenic fragments that could elicit such reactions upon binding proteins involved in the host cell cytotoxic adaptive immune response? Such a reaction was not observed in the SARS-CoV pandemic of 2003 (shortly SARS1). What is unique to SARS-CoV-2, and how recent mutations in SARS-CoV-2 S may have promoted such an increased virulence?

**Table 1:** Similarities between clinical and laboratory features of MIS-C and pediatric TSS.

| Clinical Features | MIS-C<sup>a</sup> | Pediatric TSS<sup>b</sup> |
| --- | --- | --- |
| High fever | + | + |
| Skin rash | + | + |
| Conjunctivitis | + | + |
| Oral mucosal involvement | + | + |
| Myalgia | + | + |
| Hypotension | + | + |
| Myocardial involvement (dysfunction) | + | + |
| Gastro-intestinal symptoms (vomiting, diarrhea, abdominal pain) | + | + |
| Renal involvement | + | + |
| CNS symptoms, altered mental state | + | + |
| Headache | + | + |
| High CRP | + | + |
| High Ferritin | + | + |
| High IL-6 | + | + |
| High D-dimers | + | + |
| High Procalcitonin | + | + |
| Lymphopenia | + | + |
| Reduced Platelet count | + | + |
| Increased Neutrophil count | + | + |
| Increased AST and ALST | + | + |
| High Pro-BNP | + | NA |
| High Troponin | + | NA |
| Isolation of TSS inducing bacteria (Staphylococcus or Streptococcus) | - | + |

<sup>a</sup> taken from refs (5-7); <sup>b</sup> taken from refs (8-11). + represents association with reported cases; NA: not available.

TSS can be caused by two types of superantigens (SAgs): bacterial or viral. Bacterial SAgs have been broadly studied. They include proteins secreted by *Staphylococcus aureus* and *Streptococcus pyogenes* that induce inflammatory cytokine gene induction and toxic shock. Typical examples are TSS toxin 1 (TSST1), and staphylococcal enterotoxins B (SEB) and H (SEH). They are highly potent T cell activators that can bind to MHC class II (MHCII) molecules and/or to TCRs of both CD4+ and CD8+ T cells. The ability of SAgs to bypass the antigen specificity of the TCRs results in broad activation of T cells and a cytokine storm, leading to toxic shock (12, 13). Notably SAgs do not bind the major (antigenic) peptide binding groove of MHCII, but instead bind other regions as well as the αβTCRs, directly. While early studies showed that bacterial SAgs activate T cells by binding the β-chain of dimeric TCRs at their variable domain (V) (14–16), more recent studies revealed that they can bind to either α- or β-chains, or both (17). The question is then, does SARS-CoV-2 S possess any superantigenic fragments/domains that could bind to the αβTCRs?

Here, we used computational modelling to determine whether the SARS-CoV-2 S possesses SAg activity. We demonstrate that an insert present in SARS-CoV-2 S, which is absent from SARS1 and MERS, mediates high affinity, non-specific binding to the TCR. Notably, a motif of ~20 amino acids enclosing this insert unique to SARS-CoV-2 among beta coronaviruses has sequence and structure features highly similar to those of the SEB toxin. Furthermore, our analysis indicates that a SARS-CoV-2 S mutation detected in a European strain may enhance TCR binding, suggesting such mutations may account for geographical differences in MIS-C occurrence. These finding have important implications for the management and treatment of COVID-19.

## RESULTS AND DISCUSSION

### SARS-CoV-2 spike harbors a high affinity site for TCR β-chain binding, which contains an insertion, P_681_RRA_684_, unique to SARS2

We first examined whether SARS-CoV-2 S could bind to the αβTCR. To this aim, we constructed a SARS-CoV-2 S structural model based on the cryo-EM structure resolved for the spike glycoprotein (18), and we used the X-ray structure of αβTCR resolved in a ternary complex with SEH and MHCII (17), and generated a series of structural models for possible SARS-CoV-2 S glycoprotein – TCR complex formation using ClusPro (19). Our simulations led to the binding pose presented in **Fig. 1A** as one of the most probable mechanisms of complex formation, as described in detail in Supplemental Information (*SI*) and **Supplementary Fig. S1**. Therein, the TCR binds at the interface between the S1 and S2 subunits of the spike protein, near the S1/S2 cleavage site. A closeup view of the interface between the spike and TCRVβ domain (**Fig. 1B**) reveals several strong interatomic interactions, involving residues S680-R683 on the spike, and R70-E74 and [Q52, D56] on the respective CDRs 3 and 2 on Vβ.

**Figure 1:**
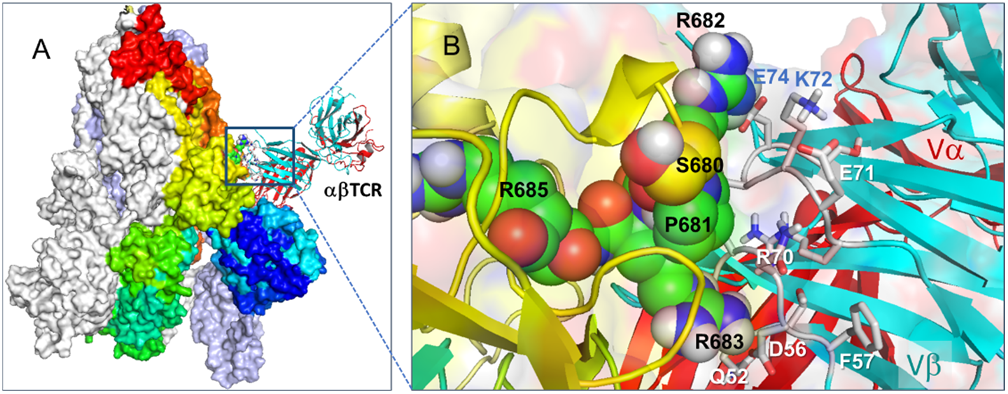
Binding of TCR to SARS-CoV-2 spike trimer near the “PRRA” insert region. Overall (**A**) and closeup (**B**) views of the complex and interfacial interactions. In (**B**) the spike monomers are colored *white*, *ice blue*, and spectrally from *blue* (N-terminal domain) to *red*, all displayed in surface representation. For better visualization, the spike trimer is oriented such that its receptor binding domains (RBDs) are at the bottom. TCR α- and β-chains are in *red* and *cyan ribbons*. In (**B**), the segment S_680_PPRAR_685_ including the PRRA insert and highly conserved cleavage site R685 is shown in van der Waals representation (*black labels*) and nearby CDR residues of the TCRVβ domain are labeled in *blue/white*. See additional information in **Supplementary Fig. S1**.

We note that the TCRVβ-binding epitope on SARS-CoV-2 S is centered around a sequence motif, P_681_RRA_684_ (or shortly **PRRA**, hereafter), and its sequential and spatial neighbors. Comparison of SARS-CoV-2 S to other beta-coronavirus spike protein sequences showed (20) that SARS-CoV-2 is distinguished by the existence of this four-residue insertion, PRRA, preceding the furin cleavage site (R685-S686 peptide bond) between the subunits S1 and S2 of each protomer (**Fig. 2A**). Structural comparison of the trimeric S proteins between SARS-CoV and SARS-CoV-2 further shows their close structural similarity in general (except for the RBD which is engaged in specific interfacial interactions (18)), but the two spikes significantly differ near the PRRARS motif unique to SARS-CoV-2, which is exposed to the extracellular medium (**Fig. 2B**).

**Figure 2:**
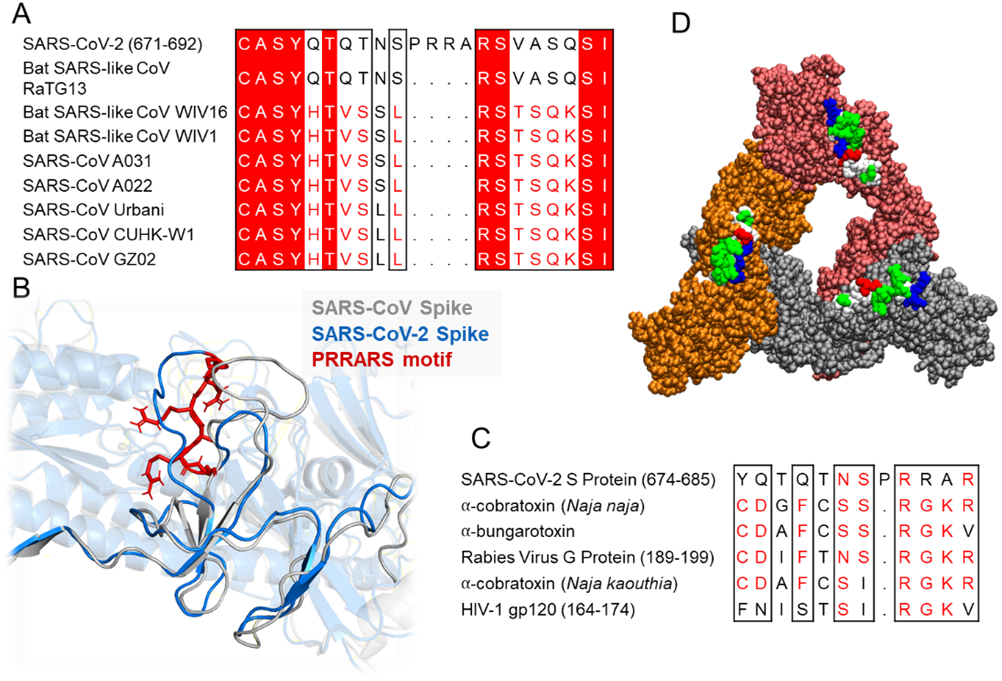
Sequence and structural properties of the insert “PRRA” motif. **A-B** SARS-CoV-2 encodes both a cleavage site (1) and neurotoxin motifs (21) near the insertion PRRA that distinguishes it from SARS-CoV. (**A**) Sequence alignment of SARS-CoV-2 and multiple SARS-CoV and Bat SARS-like CoV strains (1) near the insertion PRRA. (**B**) Structural alignment of SARS-CoV-2 and SARS-CoV at the same region. The PRRARS motif is shown in *red sticks*. (**C**) Sequence similarity between neurotoxin motifs and the close neighborhood of the PRRA insert, reported earlier (21) as well as HIV-1 gp120 SAg motif (22) in the last row. (**D**) SARS-CoV-2 S trimer composed of S1 subunits only. The protomers are colored *orange*, *red* and gray, and displayed in van der Waals format. The protruding motifs E661-R685 are highlighted in *white*, *green*, *red*, and *blue* representing the hydrophobic, hydrophilic, acidic, and basic residues.

### Further examination of the motif near PRRA reveals close structural similarity to the SEB superantigen as well as sequence similarities to neurotoxins and a viral SAg

The insertion **PRRA** together with the sequentially preceding seven amino acids and succeeding Arg (fully conserved among β-coronaviruses) have been pointed out to form a motif, Y_674_QTQTNS**PRRA**R_685_, homologous to that of neurotoxins from *Ophiophagus* (cobra) and *Bungarus genera*, as well as neurotoxin-like regions from three RABV strains (21) (**Fig. 2C**). We further noticed that the same segment bears close similarity to HIV-1 glycoprotein gp120 superantigenic motif F164-V164.

This close sequence similarity to both bacterial and viral SAgs, in support of the potential superantigenic character of the amino acid stretch Y674-R685 of SARS-CoV-2 S directed us to further analyze its local sequence and structural properties. Our analysis led to an interesting sequence similarity between the partially overlapping fragment T678-Q690 of the spike and the SEB superantigenic peptide Y_150_NKK-KATVQELD_161_ (**Fig. 3A**). This dodecapeptide sequence within the SEB shows strong conservation among a broad range of *staphylococcal* and *streptococcal* SAgs (23, 24). We note that the sequentially aligned segment of SARS1 (S664-K672) bears minimal similarity to the SEB SAg (**Fig. 3A** *left*). What is even more interesting is that SARS-Cov-2 motif showed a palindromic behavior with respect to the superantigenic SEB sequence in the sense that a broader stretch, from E661 to R685, could be aligned to the superantigen peptide in the reverse direction as well (**Fig. 3A** *right*). This brings to our attention the versatility and high propensity of the SARS-CoV-2 S TCRVβ-binding site residues to potentially act as a superantigenic fragment.

**Figure 3:**
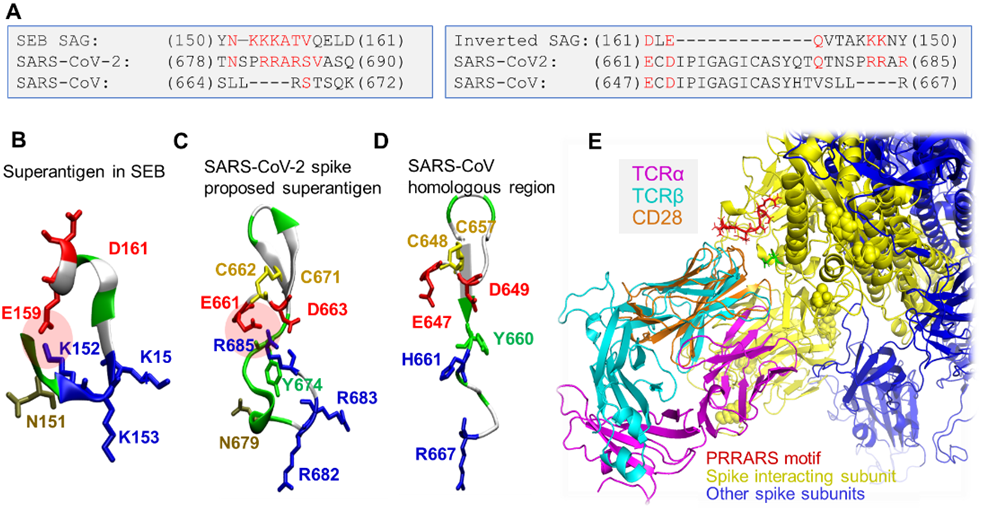
The “PRRA” insert in SARS-CoV-2 spike exhibits sequence and structure properties similar to those of bacterial superantigen SEB. (**A**) Alignment of the superantigenic sequence of SEB (23) against a homologous sequence of SARS-CoV-2 spike near the PRRA insert and corresponding SARS-CoV segment. Alignments are displayed for both forward (*left*) and reverse (*right*) ordering of the SEB sequence. Note the similarity between the former two, while the third (SARS-CoV) shows similarities to SARS-CoV-2, but not SEB, sequence. (**B**) Structure of the superantigenic peptide (T150 - D161) observed in the crystal structure of SEB (25) (PDB: 3SEB). (**C**) Structural model for SARS-CoV-2 S palindromic motif E661 - R685. (**D**) Homologous region in SARS-CoV S exhibits totally distinctive structural features: a salt bridge, K152-E159 (in SEB) or R685-E661 (SARS-CoV-2), is absent in SARS-CoV spike; the former two are poly-basic (with three lysines and three arginines in the respective motifs), whereas SARS-CoV spike counterpart has one basic residue (R667) only; and the former two possess a scaffolding ASN, which is absent on SARS1. (**E**) Structural alignment of CD28, the receptor binding SEB, onto TCRVβ domain, in support of the adaptability of the putative SAg site to accommodate spike-TCRβ or SEB-CD28 interactions.

Significantly, the structures of the two peptides exhibit a remarkable similarity (**Fig. 3B-C**), including a salt bridge stabilizing each structural motif (E159-K152 in SEB and E661-R685 in SARS-CoV-2 S), the relative orientations of three positively charged residues (K152-K153-K154 in SEB and R682-R683-R685 in SARS-CoV-2 S), a β-hairpin which apparently serves as a scaffold, and may be further supported in SARS coronavirus by a potential disulfide bond between C662-C671. The former two features are absent in SARS1 spike (**Fig. 3D**).

This analysis overall indicates that the segment **T_678_NSPRRAR**_685_ forms a putatively *superantigenic core*, consistently aligned against various bacterial or viral SAgs (**Figs. 2C** and **3A-C**) with or without the participation of the adjoining amino acids. However, combined broader sequence and structure analysis in **Fig. 3** panels **A** (*right*) and **B-C**, reveals an even more compelling feature: this putative SAg core is structurally consolidated by spatial proximity to a conserved acidic segment, E_661_CD_663_, which forms a highly stable salt bridge with the polybasic segment PRRAR of SARS-CoV-2 S, much in the same way as to the salt bridge observed in SEB (but not in SARS1 S).

We note that the SEB superantigen peptide Y_150_NKKKATVQELD_161_ has been reported to bind CD28 (23), a T cell receptor that provides co-stimulatory signals required for T cell activation and survival. CD28 and TCRV domains share the same (immunoglobulin) fold (**Fig. 3E**), and the binding mechanism shown in **Fig. 1B** is likely to be adopted with minor rearrangements for enabling the binding of SEB to CD28.

Finally, because of the homologous superantigenic segment of SEB binding CD28, we also tested the potential binding of SARS2 spike E661-R685 onto CD28, considering the possibility that the target of SARS2 spike superantigenic segment might be CD28. Our simulations indicated that the same segment can equally bind to CD28, further supporting the strong propensity of the fragment to stimulate T cell activation.

### SARS-CoV-2 spike retains other superantigenic or toxic fragments observed in SARS1 spike, including an ICAM-1 like motif engaged in stabilizing interactions with TCRVα

The existence of potential superantigenic, toxic or intercellular-adhesion molecule (ICAM)-like sequence fragments in SARS1 was thoroughly examined by Li et al. following the 2003 pandemic (26). This led to the identification of the nine sequence stretches including three *Botulinum* neurotoxin type D or G precursors, and two motifs that have a high similarity with the intercellular adhesion molecule 1 (ICAM-1). Comparative analysis with SARS-CoV-2 spike sequence revealed that seven of these sequence motifs are conserved between SARS-CoV and SARS-CoV-2 (**Supplementary Fig. S2**). Among them, we note that Y_279_NENGTITDAVDCALDPLSETKC_301_, an ICAM-1 (CD54)-like motif, also participates in the association between the SARS-CoV-2 spike and the bound αβTCR (see **Fig. 4**).

**Figure 4:**
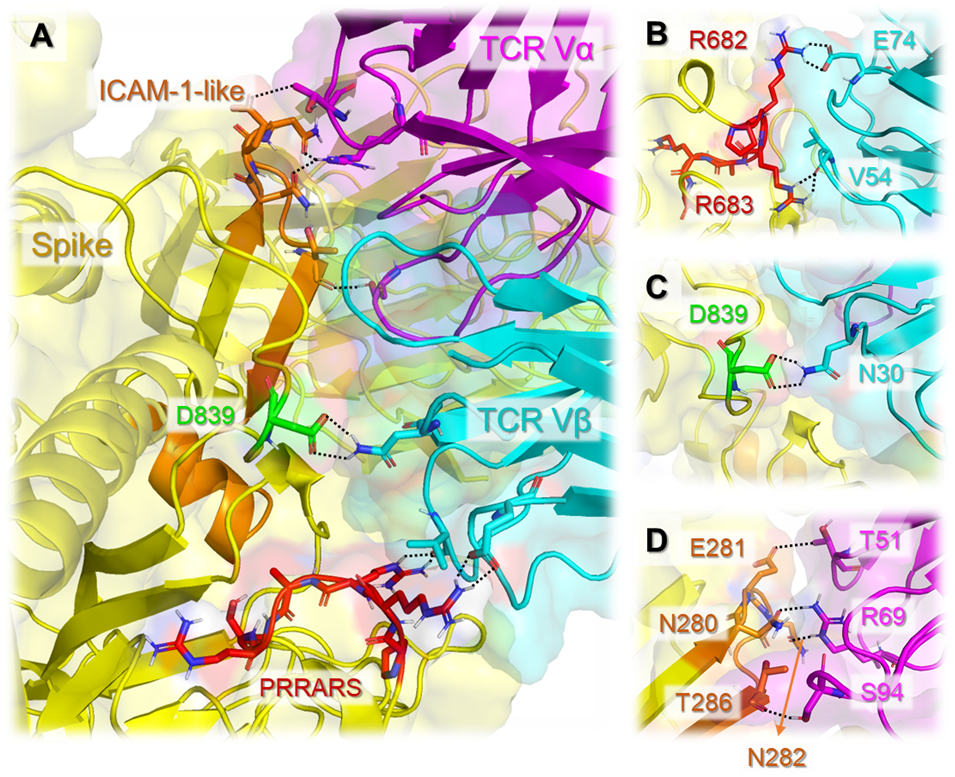
The interfacial interactions between SARS-CoV-2 spike and αβTCR are further stabilized by the association of an ICAM-like motif with TCRVα domain. (**A**) Interface between SARS-CoV-2 spike and TCR variable domains. Spike is shown in *yellow*; TCR Vα and Vβ are in *magenta* and *cyan*, respectively. The PRRARS insert is high-lighted in *red*; The mutation site D839 identified in recent study (28) is in *green*; SARS-CoV-2 counterpart of CD54-like motif identified for SARS-CoV spike (26) is in *orange*. Residues involved in close interfacial contacts are shown in *sticks*, with nitrogen and oxygen atoms colored *blue* and *red*, respectively. Interactions between atom pairs separated by less than 2.5Å are indicated by *black dashed lines*. (**B**) A close-up view of the interactions between the PRRARS insert/motif and TCR Vβ. (**C**) Same for the D839 mutation site. (**D**) Interactions between selected residues on ICAM-1-like motif (*labeled, orange*) TCRVα CDRs.

Note that ICAM-1 involvement is critical to mediating immune and inflammatory responses. The observed interaction of the ICAM-1-like motif of SARS-CoV-2 S with TCRVα, in tandem with the interaction of the above discussed putative SAg motif (around the insert PRRA) with TCRVα, is likely to further strengthen the association of the virus with the T cell and the ensuing activation. Precisely, N280-E281-N282 and T286 belonging to the ICAM-like fragment closely interact with the TCRVα CDRs; mainly T286 (spike) makes close contacts with S94 (CDR3), E281 (spike) forms a hydrogen bond with T51 (CDR2), and N280 and N282 (spike) closely associate with R69 (**Fig. 4D**).

### A rare mutation, D839Y/E, recently observed in a SARS2 strain from Europe contributes to stabilizing the interaction with TCR

Interestingly, the SARS-CoV-2 spike binding region harbors three residues that have been recently reported to have mutated in new strains from Europe and USA (27, 28): D614G, A831V and D839Y/N/E). The former two may potentially interact with MHCII; while the latter (D839, European strain) is located close to TCRVβ and strongly interacts with N30; (**Fig. 4A** and **C**, and **Supplementary Figs. S3** and **S4A**). Its substitution by glutamate in the mutant D839E increases the strength of the intermolecular (and thereby virus-T cell) association (**Supplementary Fig. S4C**). Even stronger interactions between spike and TCRVβ are observed upon replacing D839 with a tyrosine as illustrated in **Supplementary Fig. S4D**: The interfacial interactions in this case are further stabilized by a hydrogen bond between Y839 and D32; an aromatic (polar-π) interaction between Y839 and N30; as well as possible electrostatic interactions with K73 and S97. Quantitative evaluation of the change in binding affinity between the spike and TCR upon mutating D839 to Y, E and N yields ΔΔG_D→Y_ = −2.2kcal mol^−1^, ΔΔG_D→E_ = −2.1kcal mol^−1^, and ΔΔG_D→N_ = −1.3kcal mol^−1^ respectively (see **Supplementary Table S1** for details). Thus, the D839Y/N/E mutations would be expected to strengthen/support the above described association between the superantigenic PRRA-containing segment and the TCRVβ.

## CONCLUSION

An understanding of the immunopathology leading to severe manifestations of COVID-19, in both adults and children, is of critical importance for effective management and treatment of the disease. MIS-C shows remarkable similarity to pediatric TSS (5–9). Using *in silico* modeling and analysis, we found that SARS-CoV-2 encodes a superantigen motif near its S1/S2 cleavage site. This region is highly similar in structure to the SEB SAg motif that interacts with both the TCR and CD28 (23) and mediates TSS. SEB enables large-scale T cell activation and proliferation (13), resulting in massive production of pro-inflammatory cytokines including IFNγ, TNFα and IL-2 from T cells as well as IL-1 and TNFα from APCs (13). This cytokine storm leads to multi-organ tissue damage similar to what is now observed in MIS-C. We therefore propose that MIS-C observed in COVID-19 patients may be mediated by superantigen activity of the SARS-CoV-2 S protein.

To date, MIS-C is mostly observed in Europe and East Coast of North America, and has not been described in Asia, despite sizeable outbreaks of COVID-19 (5–7) (CDC and ECDC). We show that a mutation at D839 found in a European strain of SARS-CoV-2 enhances the binding affinity of the SAg motif to the TCR. This could (at least partly) explain the geographical skewing of MIS-C to areas where the European strain is endemic, and identification of other strain-specific mutations may help predict where future outbreak of MIS-C may occur.

Our findings raise the exciting possibility that immunomodulatory therapeutic options used for TSS may also be effective for MIS-C, including IVIG and steroids. Indeed, initial published and unpublished reports suggest that MIS-C patients respond well to IVIG with or with-out steroids (5–7). IVIG recognizes SEB epitopes (29), and thus may function in part by neutralization of a superantigen. Given structural similarities between SEB and the S protein SAg motif, there is potential for cross-reactivity of these immunoglobins, explaining at least in part the response of MIS-C cases to IVIG. Other FDA-approved anti-inflammatory drugs tested in models of SEB TSS may also be effective, including CTLA4-Ig which can inhibit CD28 co-stimulation (30), and the mTOR inhibitor rapamycin (31), which is already in use for COVID-19. In addition, humanized monoclonal anti-SEB Abs have been described (32) that could also be of potential therapeutic benefit in MIS-C patients. Notably, it has been shown in the mouse model of TSS that lethal SEB superantigen challenge can be prevented by short peptide mimetics of its superantigen motif (23). It would be interesting to examine whether short peptide mimetics of SARS-CoV-2 spike superantigen region might be employed to prevent/attenuate inflammatory cytokine gene induction and toxic shock in MIS-C patients.

At present, the majority of antibody therapies under investigation are designed to target the SARS-CoV-2 receptor binding domains (RBDs) (33, 34), and our simulations also indicated that RBD might potentially interact with TCRs. However, compared with RBDs, relatively fewer mutations are found in the SAg region of SARS-CoV-2; notably, the “PRRA” insert is unique to SARS-CoV-2 and retained among all of its isolates sequenced to date (27, 28). It might be constructive to design antibodies or drugs targeting this SAg region, to not only block the cleavage essential to enabling viral entry (1, 20), but also modulate the SAg-induced inflammatory cytokine gene induction (13).

Fortunately, severe respiratory manifestations of COVID-19 in children as well as development of MIS-C are rare. This may be due to trained immunity (2) or cross-viral immunity to other coronavirus strains (35). T and B cells play an important role in the anti-viral response. CD4+ and CD8+ T cells from convalescent COVID-19 patients can recognize a range of SARS2 epitopes, and the S protein represents a major target (35). Interestingly, T cells from unexposed individuals can also respond to S protein epitopes from SARS-CoV-2, which supports the hypothesis of cross-viral immunity from other coronavirus strains (35). However, why only a fraction of infected children develop MIS-C is unclear. It is possible that a poor initial antibody response to the virus fails to neutralize SAg leading to immune enhancement following re exposure or that certain HLA types are more permissive of binding SAg, and indeed HLA has been shown to play a role in COVID susceptibility (36). Of the nine cases initially reported in the UK, six were of Afro-Caribbean descent, which also suggests a potential genetic component to susceptibility (5).

It is interesting to note that approximately a third or fewer of MIS-C patients tested positive for the SARS-CoV-2, but the majority (but not all) have serologic evidence of infection or a history of exposure to COVID-19 (5–7). This may suggest that the SARS-CoV-2 SAg causes a delayed hyperinflammation response in certain children. SAgs have been implicated in autoimmunity by triggering self-reactive T cells (12). Antibody-mediated enhancement upon re-exposure to the virus may also contribute to uncontrolled infection and inflammation (37). It is also possible that despite a negative nasopharyngeal PCR test, the virus may still be present in the gastrointestinal tract (38). MIS-C patients demonstrate unusually severe GI symptoms, abdominal pain, vomiting and diarrhea, in addition to severe myocardial dysfunction and cardiac shock (5–7) and such severe GI symptoms are also frequently associated with the SAg (9).

In summary we made three major observations: (a) PRRAR and sequential neighbors interact with TCRVβ residues D56, R70 and E74 at the CDRs, and this association closely resembles that of SEB SAg with TCRVβ; (b) nearby D839 participates in this interaction and its mutation to tyrosine would further strengthen the association with TCRVβ; and (c) a sequence motif (N280-T286) typical of ICAM-1 further interacts with the TCRVα further stabilizing/enhancing the association between the viral spike and host cell TCR. So, the binding pose of TCR with respect to the spike allows simultaneously for all three associations. The first is the most important. The 2^nd^ is significant because it is the European strain. The 3^rd^ may be imparting a stronger host-virus adhesion. Therefore, strategies used for the treatment of SEB-mediated Toxic Shock or approaches to block the interaction of S protein with the TCR may reduce hyperinflammatory manifestations of COVID-19 in both adults and children.

## MATERIALS AND METHODS

SARS-CoV-2 (P0DTC2) and SARS-CoV (CVHSA_P59594) spike models were generated using SWISS-MODEL (39), based on the resolved spike glycoprotein structures of SARS-CoV-2(18) (PDB: 6VSB) and SARS-CoV (40) (PDB: 6ACD).The missing loops in the crystal structures were built using libraries of backbone fragments (41) or by constraint space *de novo* reconstruction of these backbone segments (42). Two mutants associated with European Covid-19 patients (28) were constructed using CHARMM-GUI (43): one is the main strain mutant D614G and the other contains four mutations including Q239K, A831V, D614G and D839Y. These two SARS-CoV-2 spike mutants together with the SARS-CoV-2 (P0DTC2) originally taken from Wuhan were used to investigate the binding to αβTCR, and MHCII (PDB: 2XN9) (17) using ClusPro (19) and PRODIGY (44). See details in the *SI.*

## Acknowledgments

We gratefully acknowledge support from NIH awards P41 GM103712 (to IB) and R01 AI072726 (to MA).

## Author contributions

M.H.C., M.A. and I.B. designed research; M.H.C., S.Z. and I.B. performed modeling research and analyzed data; R.A.P. and M.A. contributed clinical research and analysis; I.B. and M.A. wrote the paper with input from all authors.

