## Supplemental Materials for "An insertion unique to SARS-CoV-2 exhibits superantigenic character strengthened by recent mutations"

### Supplemental Methods

#### Generation of a binary complex between SARS-CoV-2 spike and T cell receptor (TCR)

SARS-CoV-2 spike model in the prefusion state was generated using SwissModel[1] based on the resolved cryo-EM structure (Protein Data Bank (PDB): 6VSB[2]) for the spike glycoprotein where one of the receptor binding domains (RBDs) is in the up conformation. The structure of the T cell receptor (TCR) containing both TCR$\alpha$ and TCR$\beta$ chains was taken from the crystal structure of the ternary complex between human TCR, staphylococcal enterotoxin H (SEH) and human major histocompatibility complex class II (MHCII) molecule[3]. Using protein-protein docking software ClusPro[4], we constructed *in silico* a series of binary complexes for SARS-CoV-2 spike and TCR. We obtained 30 clusters of conformations for spike-TCR binary complexes, upon clustering ~1000 models generated by ClusPro. The clusters were rank-ordered by cluster size, as recommended[4]. We analyzed all models and found that the majority (>90%) showed that TCR bound to spike via its constant domain. Given that the constant domain is proximal to the cell membrane and TCR employs the variable domain for binding superantigens and/or antigen/MHC complexes[3], we then added restraints to our docking simulations to prevent the binding of TCR constant domain and filter out those conformers where the variable domain would bind to the spike. This led to 27 clusters (based on a set of 666 models) from ClusPro. Interestingly, 45% of models showed the binding of TCR near the region of "PRRA" insert and 46% of models showed the binding of TCR within multiple RBDs. Thus, we identified two hot spots for TCR binding within SARS-CoV-2 spike: one is near "PRRA" insert and the other within the RBD. Representative members belonging to the top-ranking clusters are presented in **Fig S1**.

#### Generation of a ternary complex between SARS-CoV-2 spike, TCR, and MHCII

Structure of the human MHCII was taken from the crystal structure of the ternary complex[3] (PDB: 2XN9) between human TCR, SEH and MHCII. First, we performed docking simulations to generate binary complexes between MHCII and SARS-CoV-2 spike. Six representative MHCII-spike binary complexes were selected to explore further docking of TCR to form a ternary complex. We analyzed all predicted ternary complex models of MHCII-Spike-TCR. Potential tertiary MHCII-Spike-TCR complex models were selected following three filtering criteria: (1) TCR either binds near "PRRA" insert region or the RBD; (2) the binding regions involve homologous superantigen or toxin binding motifs predicted for SARS-CoV (**Fig. S2**); (3) MHCII and TCR are in close proximity. These filters led to the MHCII-Spike-TCR complex model illustrated in **Fig. S3A**. Interestingly, the SARS-CoV-2 spike binding region harbors three residues that have been recently reported to have mutated in new strains from Europe and USA[5,6] (**Fig. S3B**): D614G, A831V and D839Y/N/E). While we do not exclude the possible occurrence of other potential tertiary complexes, especially those involving the RBDs, we focus here on the complex shown in **Fig. S3**, which uniquely satisfied all three aforementioned criteria.

##### *In silico* mutagenesis of D839 of SARS-CoV-2 spike

We mutated D839 of the SARS-CoV-2 spike *in silico* to asparagine, glutamic acid and tyrosine in line with the aforementioned mutant D839Y/N/E observed in a new strain from Europe. To this aim, we used PyMOL mutagenesis tool[7] and evaluated the change in local conformation and energetics in the complex formed with TCR. The most probable rotamers were selected and energetically minimized in the presence of the bound TCR (conformation shown in **Fig. 1**) using OpenMM[8]. Binding affinities (ΔG) and dissociation constants ($K_d$) were obtained for (i) the full complex (with the intact spike and entire TCR as interactors) or (ii) a single spike subunit and TCRVβ with the D839Y/N/E mutation on spike at 37 °C using PRODIGY server[9,10].

#### Supplemental Figures and Tables

**Table S1: Binding affinities predicted for the interactions between the αβTCR and SARS-CoV-2 spike before/after the point mutation D839Y/N/E.**

| | Aspartic Acid (D) | | Tyrosine (Y) | | Glutamic Acid (E) | | Asparagine (N) | |
|---|---|---|---|---|---|---|---|---|
| | ΔG (kcal mol$^{-1}$) | $K_d$ (nM) | ΔG (kcal mol$^{-1}$) | $K_d$ (nM) | ΔG (kcal mol$^{-1}$) | $K_d$ (nM) | ΔG (kcal mol$^{-1}$) | $K_d$ (nM) |
| **Full complex** | -11.0 | 18 | -13.2 | 0.46 | -13.1 | 0.56 | -12.3 | 2.3 |
| **S subunit - TCR Vβ** | -8.8 | 580 | -10.3 | 53 | -10.1 | 80 | -9.5 | 190 |

* Binding affinities (ΔG) and dissociation constants ($K_d$) were obtained at 37 °C using PRODIGY server[9,10].

#### Supplemental Figures

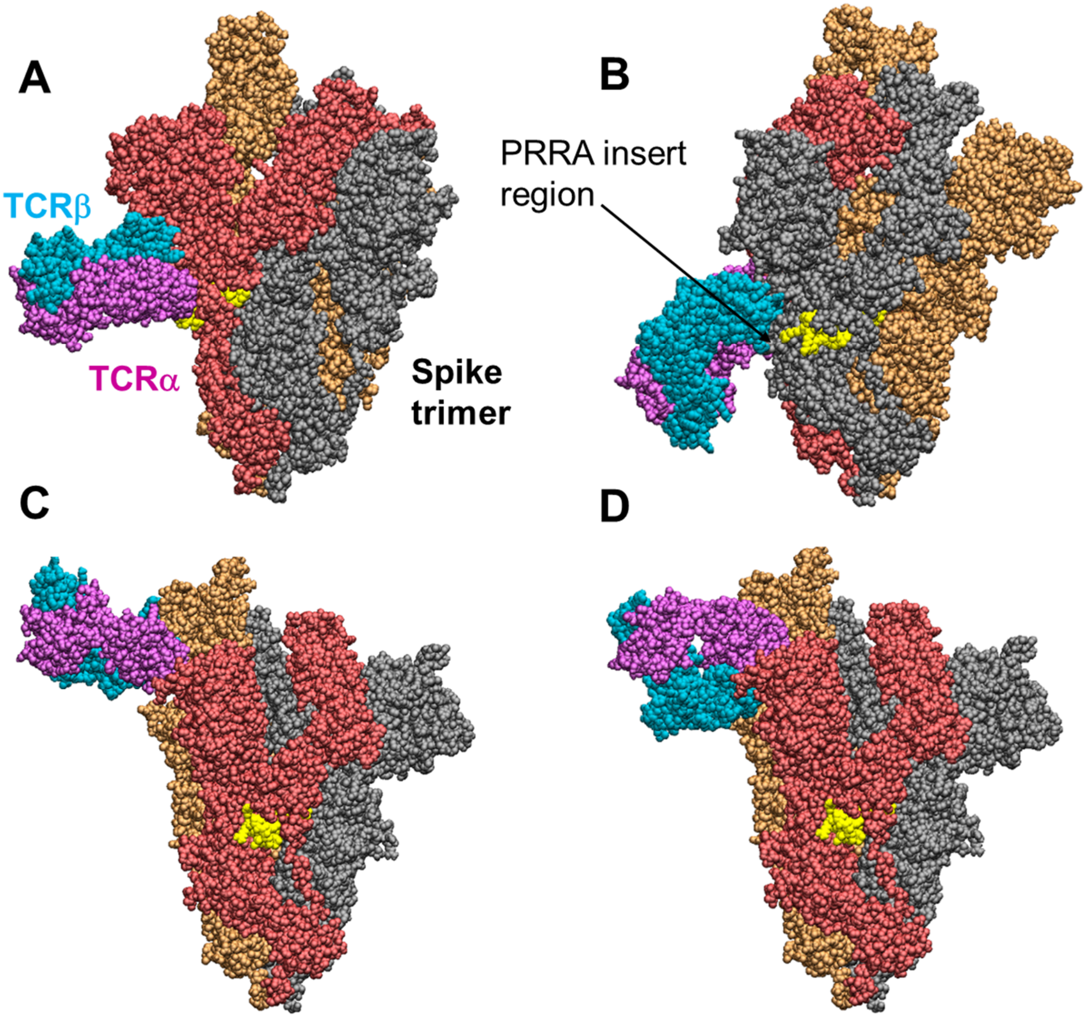


**Figure S1: Top-ranking binary complexes of SARS-CoV-2 spike with the T cell receptor (TCR) predicted by ClusPro.** (**A-B**) Binding of TCR near the "PRRA" insert region. (**C-D**) Binding of TCR near the RBD of a subunit. The spike trimer subunits are colored *red*, *orange,* and *gray*. The PRRA insert region (E661 to R685) is shown in *yellow*. TCR α- and β-chains are shown in *cyan* and *magenta*. See more details for the interaction between the PRRA insert region and TCR in **Fig. 1**.

| A | Sequence | Residues | Sequence Identity |
|---|---|---|---|
| 1 | VIPFKDGIYFAATEKSNVVRGWVFGSTM | 80-107 | 68% |
| | VLPFNDGVYFASTEKSNIIRGWIFGTTL | 83-110 | |
| 2 | QTHTMIFDNAFNCTFEYISDAFSLDVS | 147-173 | 37% |
| | ESEFRVYSSANNCTFEYVSQPFLMDLE | 154-180 | |
| 3 | NITNFRAILT---AF-SPAQDI---WGTSA | 227-249 | 30% |
| | NITRFQTLLALHRSYLTPG-DSSSGWTAGA | 234-262 | |
| 4 | YDENGTITDAVDCSQNPLAELKC | 266-288 | 74% |
| | Y**NEN**GTI**T**DAVDCALDPLSETKC | 279-301 | |
| 5 | LKCSVKSFEIDKGIYQTSNFRVVPSGDVVRFPNITNLCPFGEVFNATKFPSVY | 286-338 | 75% |
| | TKCTLKSFTVEKGIYQTSNFRVQPTESIVRFPNITNLCPFGEVFNATRFASVY | 299-351 | |
| 6 | GCLIGAEHVDTSYECDIPIG | 634-653 | 90% |
| | GCLIGAEHVNNSYECDIPIG | 648-667 | |
| 7 | NTREVFAQVKQMYKTPTLKYFGGFNFSQILP | 759-789 | 84% |
| | NTQEVFAQVKQIYKTPPIKDFGGFNFSQILP | 777-807 | |
| 8 | EAEVQIDRLITGRLQSLQTYVTQQLIRAAEIRASANLAATKMSECVLGQSKRVDFCGKGYHLMSFPQAAPHGVVF LHVTYVPS | 970-1052 | 98% |
| | EAEVQIDRLITGRLQSLQTYVTQQLIRAAEIRASANLAATKMSECVLGQSKRVDFCGKGYHLMSFPQSAPHGVVF LHVTYVPA | 988-1070 | |
| 9 | LQPELDSFKEELDKYFKNHTSPDVDLGDISGINASVVNIQKEIDRLNEVAKNLNESLIDLQ | 1123-1183 | 100% |
| | LQPELDSFKEELDKYFKNHTSPDVDLGDISGINASVVNIQKEIDRLNEVAKNLNESLIDLQ | 1141-1201 | |

B

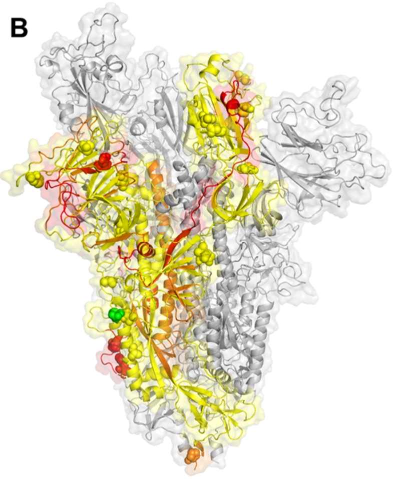

**Figure S2: Motifs associated with superantigen, toxin, cytokine, and membrane surface proteins predicted for SARS-CoV spike and mapped onto SARS-CoV-2 spike sequence and structure.** (**A**) Sequence alignment of these motifs on SARS-CoV (*upper rows*) and SARS-CoV-2 spikes *(lower rows),* corresponding residue numbers (3rd column) and sequence identity (4th/last column). Superantigenic and toxic-like motifs are highlighted in *pink*. Residues that interact with TCR Vα are marked in *red*. (**B**) Predicted motifs mapped onto the trimeric structure of SARS-CoV-2 spike, with one of its subunit colored in *yellow*. The motifs are colored *red* (superantigenic and toxic-like) or *orange* (others). Mutation sites reported in recent work[5,6] are shown in *spheres*. The mutation site D839Y/N/E is highlighted in *green*.

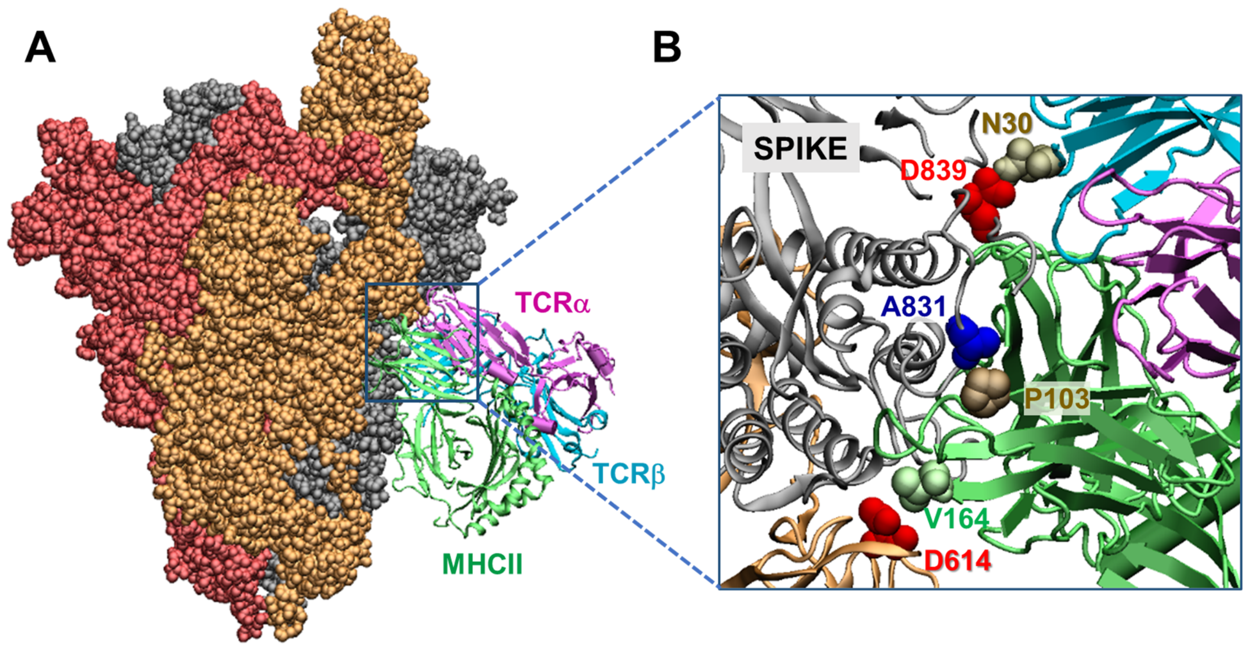


**Figure S3**: **Modeled ternary complex formed by SARS CoV-2 spike, MHCII and αβTCR**. (**A**) Side view of the complex. The trimeric spike subunits are shown in the same style and color as in **Fig. S1**, as well as TCRα and TCRβ. The TCR retains a similar pose as in panel **B** in **Fig. S1**, now rotated by 180˚ along the z-axis. MHCII is displayed in *green* ribbon diagram. (**B**) Top view of the interfacial contacts. Note that three spike residues reported to mutate in recent strains observed in European and western counties[5,6] (D614G, A831V and D839Y/N/E) are within 3-5 Å from either MHCII or TCR Vβ.

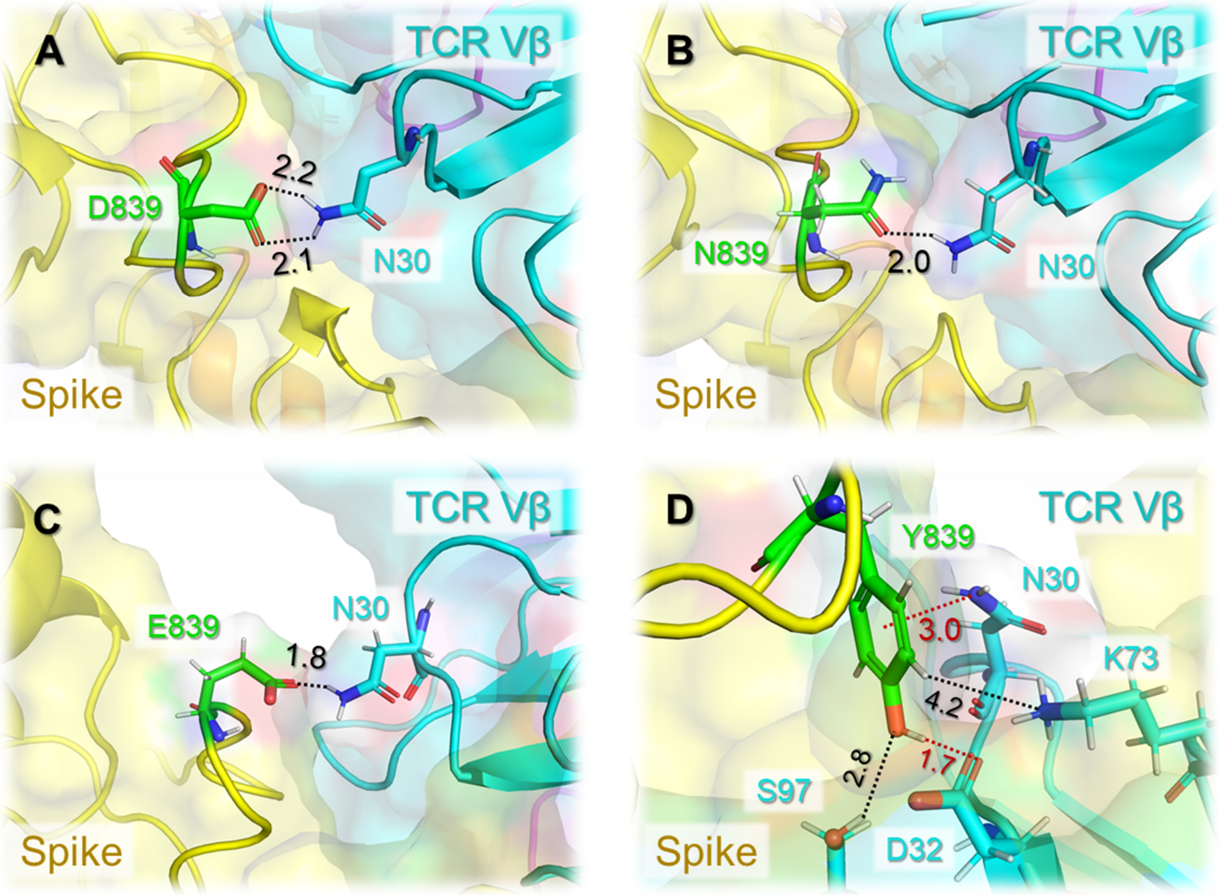


**Figure S4:** ***In silico* mutagenesis analysis of SARS-CoV-2 spike protein residue D839Y/N/E.** (**A**) A close-up view of the interaction between the wild type residue D839 in the spike and N30 of TCR Vβ. (**B-D**) Results obtained upon mutation to asparagine, glutamic acid, and tyrosine. The spike and TCR Vβ are shown in *yellow* and *cyan*, respectively. The mutation site is highlighted in *green*. Atomic interactions are indicated by *black dashed lines* along with their distances in Ångstroms.
